# DNA metabarcoding diet analysis reveals dynamic feeding behaviour and biological control potential of carabid farmland communities

**DOI:** 10.1101/332312

**Authors:** Stefaniya Kamenova, Vincent Bretagnolle, Manuel Plantegenest, Elsa Canard

## Abstract

Maximizing the delivery of key ecosystem services such as biological control through the management of natural enemy communities is one of the major challenges for modern agriculture. The main obstacle lies in our yet limited capacity of identifying the factors that drive the dynamics of trophic interactions within multi-species assemblages. Invertebrate generalist predators like carabid beetles are known for their dynamic feeding behaviour. Yet, at what extent different carabid species contribute to the regulation of animal and plant pests within agroecosystems is currently unknown. Here, we developed a DNA metabarcoding approach for characterizing the full diet spectrum of a community of fourteen very common carabid species inhabiting an intensively managed Western-European agroecosystem. We then investigated how diet and biological control potential within the carabid community varies with the sampling field location and the crop type (wheat *vs* oilseed rape). DNA metabarcoding diet analysis allowed to detect a wide variety of animal and plant taxa from carabid gut contents thus confirming their generalist feeding behaviour. The most common prey categories detected were arachnids, insects, earthworms and several plant families potentially including many weed species. Our results also show that the field location and the crop type are much stronger determinants then the species regarding carabid dietary choice: significantly more trophic links involving dipteran prey were observed in wheat, whereas more collembolan and plant prey was consumed in oilseed rape by the same carabid community. We speculate that structural differences in the habitats provided by these two crop types drive differences in resource availability cascading up the trophic chain, and we assume that specific carabid taxa could hardly be used to infer levels of ecosystem services (biological control) or disservices (e.g. intraguild predation). However, as this is the first study to report the use of DNA metabarcoding diet analysis in predatory carabid beetles we urge caution over the interpretation of our results. For instance, overall detection rates were rather low (31% of the individuals analysed tested positive for at least one prey category) most likely due to the overwhelming amplification of the carabid host DNA. Therefore, we acknowledge that more studies are required in order to confirm our observations and conclude with few recommendations for further improvements of the community-level DNA metabarcoding analysis of carabid diet.

## Introduction

Agroecological intensification has been proposed as a viable alternative to conventional crop management that relies on large agrochemical input for maintaining yield levels (Tscharntke et al. 2012). One of the main premises of agroecology is that sustainable agriculture and global food security could only be guaranteed by incorporating into the management agenda all the ecosystem services provided by biodiversity that naturally enhance crop production (Bommarco et al. 2013). Indeed, in biodiversity-friendly cropping systems, important ecological processes such as nutrient recycling or beneficial trophic interactions (e.g. pollination, pest predation) are usually preserved and enhanced by high levels of functional biodiversity (Moonen & Bàrberi 2008; Mace et al. 2012). Biological control is typically an important ecosystem service that results from the direct and indirect trophic interactions linking pest species and natural enemies guilds within agricultural areas. However, predicting when and why these trophic interactions provide an efficient pest regulation still remains a daunting challenge for both scientists and stakeholders. Previous studies have shown that an increase in functional diversity of natural enemies does indeed strengthen pest regulation (Losey & Denno 1998; Letourneau et al. 2009), but compiling evidence also shows that species-rich predator assemblages could have neutral or even negative effects on pest suppression (Finke & Denno 2004; Martin et al. 2013). Such context-dependency suggests that beyond the diversity of natural enemies guilds, additional factors drive pest-enemies interactions. Ecosystem services such as biological control typically emerge from the complex network of trophic interactions among multiple species at multiple trophic levels-a complexity that may often obscure the exact mechanisms behind the relationship between biodiversity and biological control in agroecosystems (Mace et al. 2012). As a consequence, there has been an increasing solicitation in the scientific community for endorsing a more mechanistic approach in the studies tackling biological control efficiency. A major obstacle in such studies lies in the difficulty to elucidate and quantify trophic interactions in highly diversified arthropod communities in the field. Recently, studies have started to examine how variations in functional traits could mediate ecological processes such as pest suppression. Results show that species functional traits appear as good predictors of both predation rates and occurrence of antagonistic interactions such as intraguild predation (Rouabah et al. 2014; Rusch et al. 2015; Brousseau et al. 2017). However, conclusions from these studies are mainly based on observations of a limited set of species or feeding interactions in laboratory conditions making it unclear how applicable these results would be in a multi-trophic context. Besides, the link between functional attributes and feeding behaviour is not always consistent, especially for generalist predators, which usually display very dynamic feeding behavior as a response to the frequent variations in prey abundance within agricultural fields (e.g. Bohan et al. 2000; Bell et al. 2010). Another way to get a mechanistic insight about processes behind pest suppression is to directly record trophic interactions within multi-species assemblages in field conditions. Instead of correlating functional trait values with pest population dynamics, a multi-trophic approach allows to directly estimating, for instance, the degree of trophic complementarity or trophic antagonisms (e.g. competition, intraguild predation) between species while directly quantifying pest consumption. Yet, such approach requires significant capacity to simultaneously characterize the diet of multiple species at various trophic levels. Relying on traditional methods such as macroscopic identifications of prey remains could be extremely laborious or even impossible in the case of liquid feeders such as most of the generalist farmland arthropod predators are (e.g. spiders, carabid beetles). DNA metabarcoding approach recently emerged as a valuable alternative for the direct characterization of complex mixtures of highly degraded DNA and has been successfully used for the diet analysis of various organisms (Ibanez et al. 2013; Mollot et al. 2014; Lefort et al. 2017). The main advantage of DNA metabarcoding is that it enables the diet analysis of generalist species without imposing the need of a strong *a priori* knowledge about species diet spectrum. Moreover, the high-throughput and the continuously decreasing costs of DNA metabarcoding significantly facilitate the rapid processing of a large number of species and individuals at once. Considering these advantages, we developed a DNA metabarcoding approach for characterizing the diet within an entire community of carabid beetles in an intensively managed European agricultural landscape. Carabid beetles (Coleoptera: Carabidae) are one of the most abundant and species-rich guilds of generalist predators within agroecosystems (Kromp 1999; McCravy & Lundgren 2011). A typical carabid community encompasses several trophic levels (herbivores, carnivores) each comprising species with various degrees of trophic specialization. This implies that depending on its specific composition, the whole community has the potential to consume a large variety of animal and plant prey including pests and other beneficial organisms, thus making predictions about carabid contribution to biological control challenging. All this makes carabid beetles an ideal model for addressing the longstanding question about the factors determining the biological control potential of generalist predators within agro-ecosystems. By directly quantifying trophic interactions from field-collected individuals, we specifically ask: (i) which carabid species have the potential to contribute to biological control? (ii) is carabid species potential contribution to biological control conditional upon environmental factors such as the sampling field location or the crop type? As a corollary of (ii) we also ask whether or not prospective changes on biological control potential in different environmental contexts are brought by changes in the diet of some species. We hypothesize that carabid species do not equally contribute to biological control, as pest consumption depends on carabid species identity regardless of the local environment. However, we also hypothesize that the local environment through the diversity of crop types and cropping practices can modulate pest consumption by carabid species through notably the availability of alternative resources. To test these hypotheses, we sampled a community of fourteen carabid species occurring simultaneously in wheat and oilseed rape fields within the Long-Term Research area ‘Zone Atelier Armorique’ in Brittany, France. For minimizing the effect of temporal changes in resource availability all carabid species were sampled at a single date. For disentangling the effect of species identity from the effect of the local environment, the same carabid species were analysed in all sampling locations or crop types.

## Materials and methods

### Sampling protocol and samples processing

The Long-Term Research area ‘Zone Atelier Armorique’ is situated in the south of the Mont St-Michel Bay, Brittany, France (48° 36’ N, 1° 32’ W). ‘Zone Atelier Armorique’ is embedded within a typical Western European agricultural landscape, characterized by a mosaic of intensively managed field crops, pastures and semi-natural elements such as hedgerows (https://osur.univ-rennes1.fr/za-armorique/). Within the zone, we randomly selected three pairs of adjacent wheat-oilseed rape fields. For maximizing the retrieval of carabid species within a timeframe of 24 h, we set up a regular grid of about 50 dry pitfall traps (H = 120 mm, Ø 8.5 cm) within each field. All pitfall traps were protected from sun and rain with an opaque lid and filled with clay beads in order to prevent predatory interactions between individuals within the trap. Seven 24h-trapping sessions were carried out between April and May 2013. Beetles were collected alive and freeze-killed at-20°C directly in the pitfall traps as soon as possible after field collection (in all cases within not more than 5h after collection). Frozen beetles were sorted out rapidly for avoiding defrosting and identified to the species level according to Roger et al. (2012). Carabid abundance and species richness in each sampled field were compared among the seven collection dates and the sampling date exhibiting the highest values was selected. Among the individuals captured at this date and for each field, up to 15 individuals from the 15 most abundant species were randomly selected. When less than 15 individuals had been collected per field and per species, all available individuals were analysed. During subsampling, we maximized the retrieval of carabid individuals from the largest possible number of pitfall traps within each field. In order to prevent any contamination by environmental DNA, the carapaces of all the selected beetles were decontaminated using the cleaning procedure described by Greenstone et al. (2013). Decontaminated individuals were then dissected and gut contents placed in sterile 2-ml micro-centrifuge tubes at-20° prior molecular analyses. During dissections, forceps were flame-sterilized and the workbench was cleaned with DNA AWAY^™^ (Thermo Scientific, USA) between each dissection.

### Reference database

We set up a sequence reference database for the most common animal prey taxa encountered in our sampled fields. For this, we took advantage of (i) the arthropod specimens accidentally trapped alongside the carabid beetles within the pitfalls; (ii) on-purpose field sampling sessions for collecting the most common mollusk species in our fields. All specimens were preserved in 96° ethanol and if possible, further identified at the finest taxonomic level. The list of the referenced taxa and their taxonomic identifications could be found in Appendix S1 (Supporting information). DNA was extracted from each specimen individually. For arthropods, we use a protocol aiming at preserving the general morphology of the specimen, while small pieces of tissue were cut off for mollusks. Total DNA was extracted using the DNeasy Blood and Tissue kit (Qiagen, Germany) following the manufacturer’s instructions. All animal specimens were amplified for the long COI fragment using M13-tailed COI primer cocktail prepared by pooling an equal volume of 10 μM of five COI primers (cf Table S1, Supporting information). PCRs were carried out in a total volume of 25 μl containing 0.625 U of HotStarTaq plus DNA polymerase (Qiagen), 2 mM MgCl_2_, 0.1 μM of each dNTP, 0.2 μM of each primer, and 2 μl of arthropod DNA extract. After an initial activation of the DNA polymerase for 2 min at 94°C, the amplification was performed with 5 cycles of 30 sec at 94°C, 40 sec at 45°C, and 1 min at 72°C; followed by 35 cycles of 30 sec at 94°C, 40 sec at 51°C, and 1 min at 72°C; and a final extension of 10 min at 72°C. The same specimens were also amplified with 16SMAV – F / 16SMAV-R primers in a total volume of 25 μl containing 1 U of HotStarTaq plus DNA polymerase (Qiagen), 2 mM MgCl_2_, 0.2 μM of each dNTP, 0.2 μM of each primer, and 5 μl of DNA template. Plants were amplified using the g/h primers in a final volume of 25 μl containing 1 U of HotStarTaq plus DNA polymerase (Qiagen), 2 mM MgCl_2_, 0.2 μM of each dNTP, 0.2 μM of each primer, and 5 μl of DNA template. PCR cycling conditions for the 16SMAV – F / 16SMAV-R and g / h primers are described below. Amplicons were sequenced using the Sanger method on both strands and for each sample with the ABI3730XL analyser (Applied Biosystems) at the Génoscope, France (http://ig.cea.fr/drf/ig/Pages/Genoscope.aspx). Sequences were assembled and aligned with CodonCode Aligner V1.5.2 (CodonCode corporation, Dedham, MA, USA). Taxonomic assignments were made with the Barcode of Life Data System Identification System (IDS) for COI (www.barcodinglife.org). Reference sequences were deposited on GenBank under XXXX.

### DNA Metabarcoding diet analysis

DNA from gut contents was extracted using the DNAeasy Blood & Tissue Kit (Qiagen, Germany) according to manufacturer’s instructions. Negative controls (water instead of DNA) were included in each batch of 64 samples during DNA extractions. PCR amplifications were realized in a final volume of 25 μl using 5 μL of DNA extract as template. The mixture contained 1 U of GoTaq^®^ Flexi DNA Polymerase (Promega, USA), 2 mM MgCl2, 0.25 mM of each dNTPs, 250 μg / mL of bovine serum albumin (BSA; Sigma, USA), 0.2 μM of each primer (Sigma, USA) and finally UHQ water to bring each sample to the final volume. PCR negative controls (water instead of DNA) were run within each batch of 89 samples. All negative controls were sequenced to check for DNA contaminations. We combined four primer sets covering the full spectrum of prey taxa consumed by the carabids (Table 1). We also used a blocking oligonucleotide specific to mammalian sequences for the 16S MAV marker in order to prevent amplifications of human DNA (De Barba et al. 2014). All samples were individually tagged using a system of 36 octamers with at least five differences among them (Coissac 2012). Tags were added on the 5’-end of each forward and reverse primer in order to obtain unique tag combinations for any given PCR product. These unique tag combinations were used afterwards to assign the high-throughput sequence data to samples using the bioinformatic pipeline OBITools (http://metabarcoding.org/obitools/doc/welcome.h™l, see below for more details about the bioinformatic analyses). PCR cycling conditions for each primer set were respected as specified in the corresponding papers (Table 1).

**Table 1.**
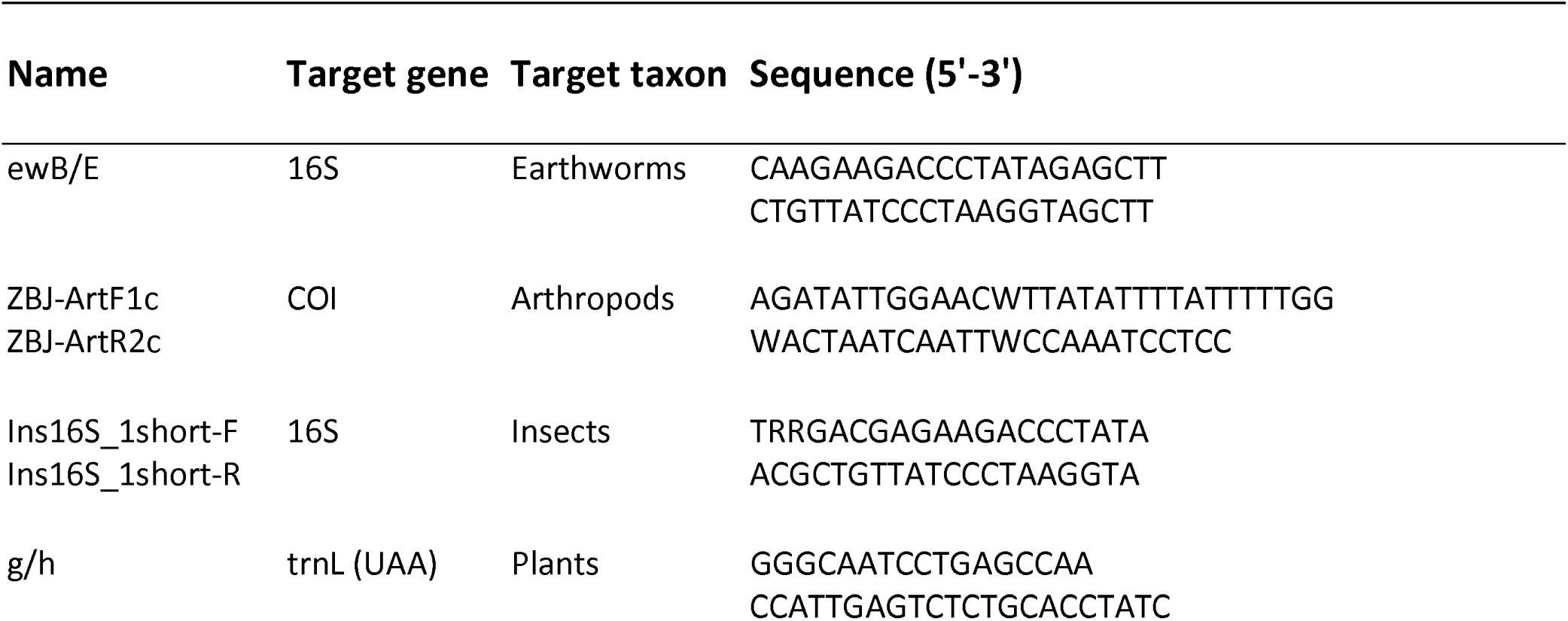
List and characteristics of the primer sets used for the DNA metabarcoding analysis of carabid diet.

All PCR products were visualized using 1.5 % agarose gel electrophoresis. According to the signal intensity of each PCR product (null, low, medium and strong, cf Mollot et al. 2014), amplicons were pooled in equimolar concentrations. Amplicons were first pooled for each primer set (n = 4), and each pool was purified using the QIAquick Gel Extraction Kit (Qiagen, Germany). Purified DNA for each primer set was quantified using Qubit fluorometer (Thermo Scientific, USA), and pooled again in equimolar concentrations resulting in one single sample sent for sequencing. Library preparation and high-throughput sequencing were carried out by Fasteris (Geneva, Switzerland). Library was prepared using the MetaFast protocol (https://www.fasteris.com/dna/?q=content/metafast-protocol-amplicon-metagenomic-analysis). The pair-end sequencing (2 × 250 bp) was carried out using an Illumina MiSeq sequencer using the Pair-end MiSeq Reagent Kit V2 following the manufacturer’s instructions.

### Bioinformatic analyses

Raw output sequences were analysed with the OBITools pipeline. First, we used the *illuminapairend* function for assembling, for each read, the forward and the reverse ends of the pair-end sequencing in one single consensus sequence (threshold quality score: ≥40). Second, consensus sequences were assigned to samples by identifying the forward and reverse primers and tag combinations using the *ngsfilter* function. All sequences that did not match perfectly with tag sequences (0 mismatch) or that had more than three mismatches on primer sequences were discarded. Third, all strictly identical sequences were clustered together (information about their distribution among samples was kept) using the *obiuniq* function. Forth, using the *obigrep* function we discarded all sequences shorter than 10 base pairs (bp) as well as all sequences occurring less than once over the entire dataset (i.e. singletons). Fifth, using the *obiclean* function that models the production of errors during the PCR (cf Boyer et al. 2016), we removed all sequences prospectively resulting from PCR errors according to the procedure described in Giguet-Covex et al. (2014). Finally, all curated sequences were taxonomically assigned using the *ecoTag* program and the EMBL sequence reference database (release 123; http://www.embl.de/). A unique taxon was assigned to each sequence. When several matches between the query sequence and the reference database were possible, the sequence was assigned to the taxon corresponding to the last common ancestor node of all the taxa in the NCBI taxonomic tree that best matched against the query sequence. A species name was accepted only if the identity score strictly equaled 1.00, a genus name in cases where the best match was ≥ 0.98, and a family name if the maximum identity was ≥0.95.

### Food web construction and statistical analyses

A carabid individual was considered positive for a prey DNA when at least one read for at least one prey taxon were sequenced. The number of sequence counts for a given prey taxon was converted into binary information (presence / absence) and the trophic links were quantified by the number of carabid individuals among the population that were positive for each prey taxon. For simplifying further analyses, prey taxa identified at various taxonomical levels were grouped into six broad resource categories: Plant, Arachnida, Clittelata, Diptera, Coleoptera and Collembola. In order to compare carabid diets between wheat and oilseed rape fields, we built bipartite food webs with the totality of the trophic links recorded. We also built bipartite food webs restrained only to what we qualified as strong trophic links-i.e. involving only carabid species for which more than 3 positive links were detected, For characterizing differences between the wheat and oilseed rape food webs, we calculated two dissimilarity indexes (trophic beta diversities, Poisot et al 2014). The first index, the whole network trophic dissimilarity, β_WN_, takes into account the totality of the links within the network (β_WN_ = 0 when the two networks share exactly the same links, β_WN_ = 1 when the two networks have no links in common). The second index, the overlapping species trophic dissimilarity, β_OS_, takes into account only the trophic links exhibited by species that are common between two networks (β_OS_ = 0 when the species exhibit the same trophic links in the two networks, β_OS_ = 1 when all trophic links differ between the two networks). For this analysis, we only considered the carabid species for which at least three individuals were positive to at least one prey category in each crop type. Dissimilarity indexes between wheat and oilseed rape food webs were compared to their random expectations based on 500 permutations of carabid individuals. First, in order to disentangle the effect of the unbalanced distribution of total number of positive carabids between wheat and oilseed rape, we permuted all individuals of all species while keeping the total number of positive individuals in each crop type (type 1). Second, in order to disentangle the effect of the differences in community composition of positive individuals between wheat and oilseed rape, we only permuted individuals for the species that were common in both crop types while keeping their local abundances constant (type 2). We used this information to deduce differences in the food web that were only due to crop plant.

Using a General Linear Model (GLM) for each resource category, we tested whether the probability of consumption of a resource by an individual was significantly influenced by (i) the sampling field location, (ii) the crop plant, (iii) the species to which it belongs:

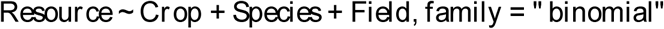

with 2 modalities for the crop plant, 14 for the carabid species and 3 for the field location. We run a best model selection using the function ‘step’ (option ‘both direction’) of the R package ‘stats’ (https://stat.ethz.ch/R-manual/R-devel/library/stats/h™l/stats-package.h™l). Because of our unbalanced data among crop types and field locations (i.e. some species being represented by more individuals than others), we also conducted a sensibility analysis by running again the model selection (as explained earlier) several times, with one particular species removed at each time.

## Results

### Carabid community in wheat and oilseed rape fields

The highest numbers of carabid species and individuals were collected on May 23^rd^. Species richness and evenness were higher in oilseed rape fields compared to wheat. Among the 47 species identified at this date, 13 major species occurred in both crop types (Fig. 1): Amara similata (AS), Brachinus slopeta (BS), Nebria salina (NS), Amara aenea (AA), Anisodactylus binotatus (AB), Asaphidion flavipes (AF), Phyla obtusa (PO), Ocydromus tetracolus (OT), Anchomenus dorsalis (AD), Metallina lampros (ML), Poecilus cupreus (PC), Loricera pilicornis (LP), and Agonum muelleri (AM). Two indivdiuals of *Laemostenus terricola* were hand-collected in oilseed rape outside the pitfall traps and included into the molecular analyses. At this date, the dominant species were *Anchomenus dorsalis, Metallina lampros* and *Poecilus cupreus* in wheat, and *Amara similata* and *Amara aenea* in oilseed rape.

**Figure 1.**
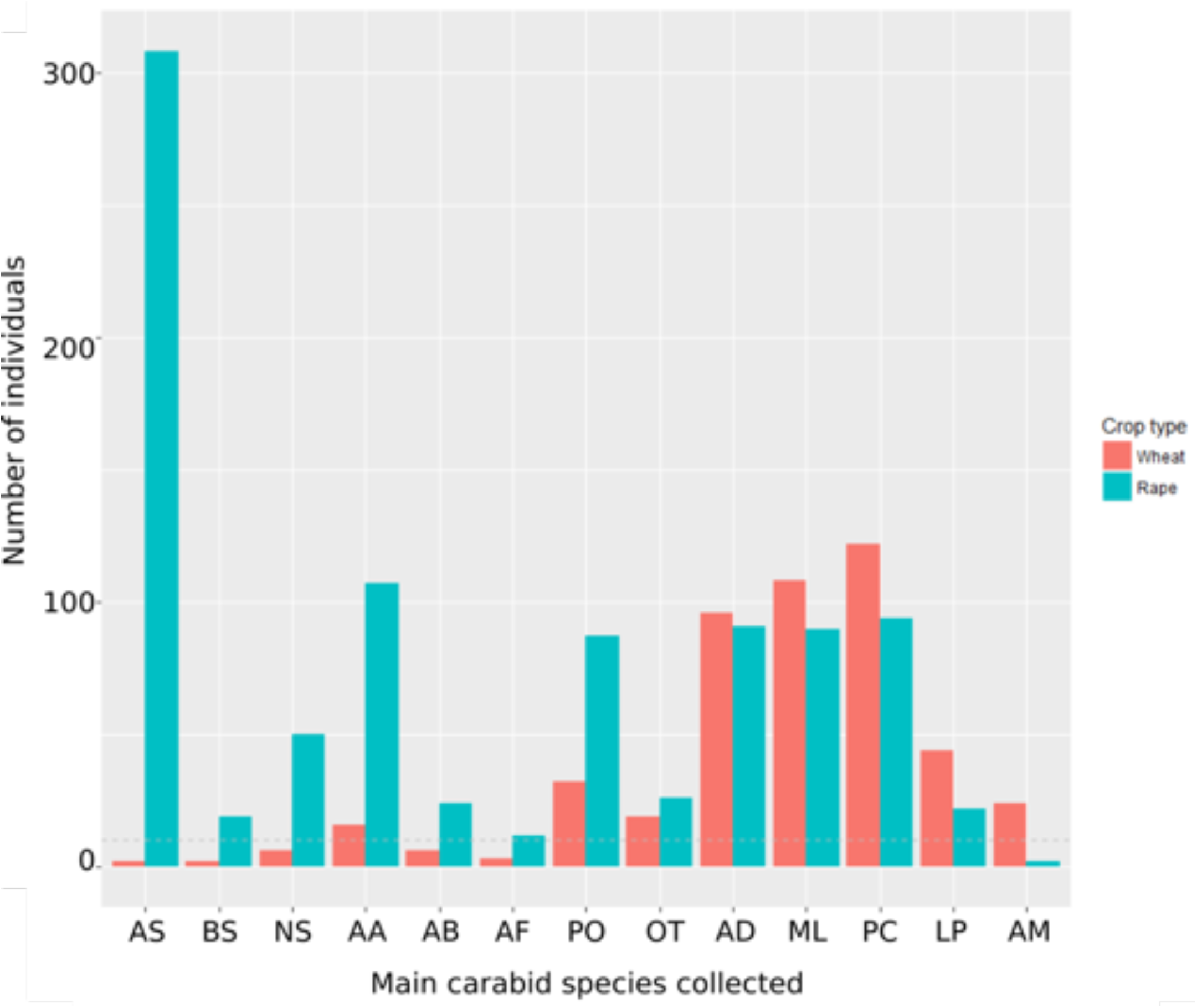
Distribution of occurrences for the most common carabid species in wheat and oilseed rape crops. Abundance of the main carabid species collected in wheat (red) and oilseed rape (green) crops on the May 23th. Abbreviations are as follows: *Amara similata* (AS), *Brachinus slopeta* (BS), *Nebria salina* (NS), *Amara aenea* (AA), *Anisodactylus binotatus* (AB), *Asaphidion flavipes* (AF), *Phyla obtusa* (PO), *Ocydromus tetracolus* (OT), *Anchomenus dorsalis* (AD), *Metallina lampros* (ML), *Poecilus cupreus* (PC), *Loricera pilicornis* (LP), and *Agonum muëlleri* (AM). Two indivdiuals of *Laemostenus terricola* (not represented here) were hand-collected outside the pitfall traps and included into the molecular analyses.

### Reference database

We successfully sequenced 291 specimens of which 265 were taxonomically assigned at least to the order level with 96% being assigned to the species level (cf Appendix S1). References specimens encompassed 8 orders of arachnids, insects and mollusks. We didn’t have any prior taxonomic reference for the mollusk specimens based on morphological identifications. This was also the case for 22 arachnid and 5 coleopteran specimens. We therefore considered that the taxonomic assignment of their corresponding barcode sequences captured their true taxonomic identity. For arachnids, we observed a mismatch between molecular and morphological taxonomic assignments for four specimens at the species level, and for one specimen at the genus level, which corresponds to 6% of all referenced specimens. For carabids this proportion was much higher with 39% of the specimens being assigned to a different genus with the molecular and morphological methods. Additional 3% of carabids were identified as different species (but the same genus) between the two methods. No mismatches were observed for any of the other referenced orders.

### DNA Metabarcoding diet analysis

74% of the sequence reads and 76% of the OTUs were assigned to the Carabidae family, which we discarded assuming they belonged to the predators themselves. 3% of the sequences reads (2% of the OTUs) were assigned to taxonomic levels that didn’t allow distinguishing prey and non-prey taxa (e.g. Hexapoda). These sequences were also discarded. Finally, 3706 sequences of 17 taxa were also retrieved from the DNA extraction and PCR negative controls, representing 0.3% of the total number of sequence reads after quality control and filtering (i.e. 1, 404 624). 93% of these contaminant sequences corresponded to Carabidae. The remaining 7% matched Plants (mainly Poaceae), Diptera (identified only as Brachicera) and Bacteria (Clostridiaceae, Clostridiaceae). The sequences of these taxa were also discarded prior statistical analyses. The reads from two OTUs both matching *Allolobophora chlorotica* with 100% identity after BLAST assignment were merged together. The remaining 52 OTUs matched prey DNA corresponding to a total of 37 animal and plant taxa that we grouped into the following six resource categories: Earthworms (4 taxa), Arachnids (6 taxa), Coleopterans other that carabids (2 taxa), Dipterans (12 taxa), Collembolans (2 taxa) and Plants (27 taxa). Prey OTUs amplified with different primers (i.e. COI and 16S MAV) were considered as separate taxa even when they matched the same species or resource categories. Among the 496 carabid individuals analysed, only 154 were positive to at least one prey DNA that occurred in >10 reads, with detection rates being higher in oilseed rape compared to wheat (19% vs 12%).

For animal prey taxa, 10 OTUs were identified to the species and 7 to the genus level allowing us to categorize them either as pests or as beneficial organisms. For plant taxa, only one OTU was identified at the species level but 11 others were identified to the genus (*Medicago* spp, *Trifolium* spp. etc.). Some of the remaining plant OTUs matched plant families containing weed species (e.g. Papaveraceae, Asteraceae, etc.). Six plant OTUs (3075 reads) were assigned to tree taxa (e.g. Fagaceae, Salicaceae, etc.). We decided to keep these taxa into further analyses as they still may result from the ingestion of tree pollen during feeding (intentionally or accidentally through the consumption of a prey covered with pollen, etc.).

### Carabid-prey food webs

Using the trophic data described above, we built bipartite food webs between carabids and their prey in wheat or in oilseed rape crops (Fig. 2). Trophic dissimilarity indexes suggested important differences between observed wheat and oilseed rape food webs (β_WN_ > 0.8, red line in Fig. 3). According to our permutation analysis, network dissimilarity between crop types remained significantly higher than expected even when differences in carabid community composition was taken into account (Fig. 3). The bipartite food webs restrained only to the strong trophic links (Fig. 4) allowed us to compare carabid species diets between wheat and oilseed rape fields. We observed than individuals of the same species could exhibit different diets in wheat and oilseed rape crops. We also observed significantly more trophic links involving dipteran prey in wheat crops, whereas more collembolan and plant prey were consumed in oilseed rape. This is coherent with the strong trophic dissimilarity indexes values (Fig. 3, β_OS_ > 0.8, red line in Fig. 3). However, carabid species factor was not retained in our best models suggesting that our data do not show strong association between carabid species and particular resource categories (Table 2). With the sensibility analysis the same best model was selected whether only species in common between the two crop types were included (i.e excluding *A. muelleri, B. sclopeta,* and *L. terricola*) or whether one species at a time was removed (Table S2, Supporting information). Results were particularly consistent for the coleopteran, arachnid and plant prey categories, with the same significance levels and very similar coefficients between the two GLMs (Table S2, Fig. S1, Supporting information). For Diptera, only the effect of the field location was marginally significant and became non-significant when removing *A. aenea, A. binotatus, A. flavipes* or *A. similata* carabid species (Table S2, Supporting information). Earthworms and Collembola prey categories were also sensible to individual species removing. Results for earthworms were always non-significant when removing any of the carabid species, while Collembola was only sensitive to the removal of the *P. cupreus* species.

**Table 2.**
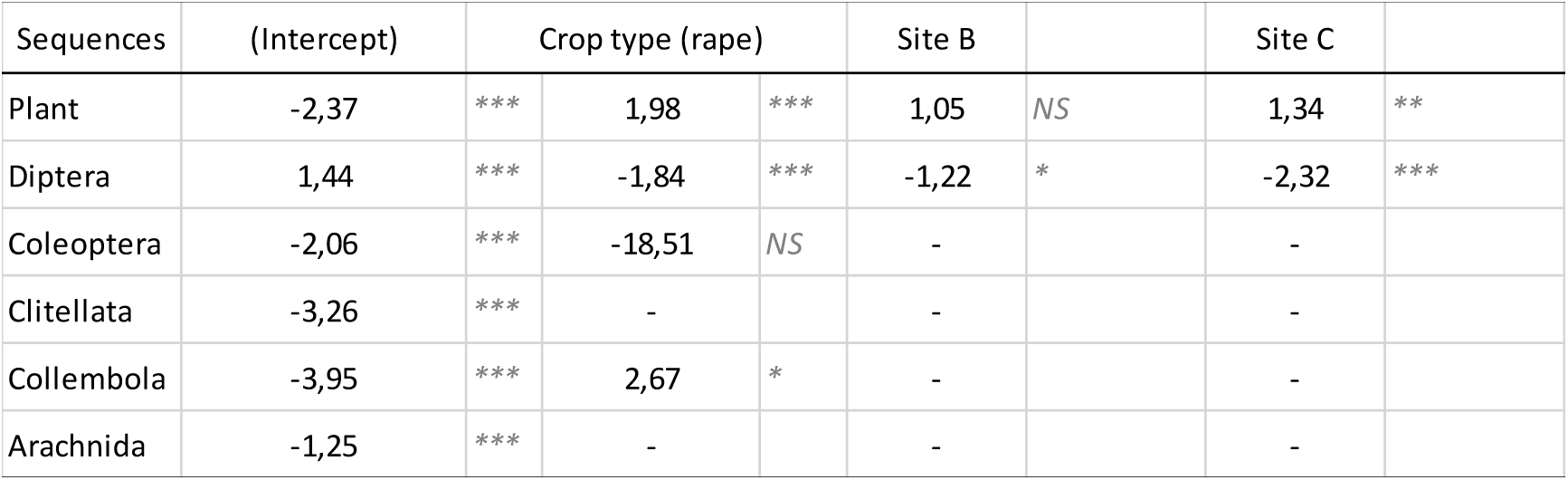
Best GLM models of resources consumption. Generalized Linear models of OTU presence / absence in carabids individuals in function of crop types, sites, and species. To perform the glm models, we retain only species that are represented by at least 3 positive individuals (e.g. 10 of the 14 species). NS: p-value > 0.05; * : p-value < 0.05; ** : p-value < 0.01; *** : p-value < 0.001.

**Figure 2.**
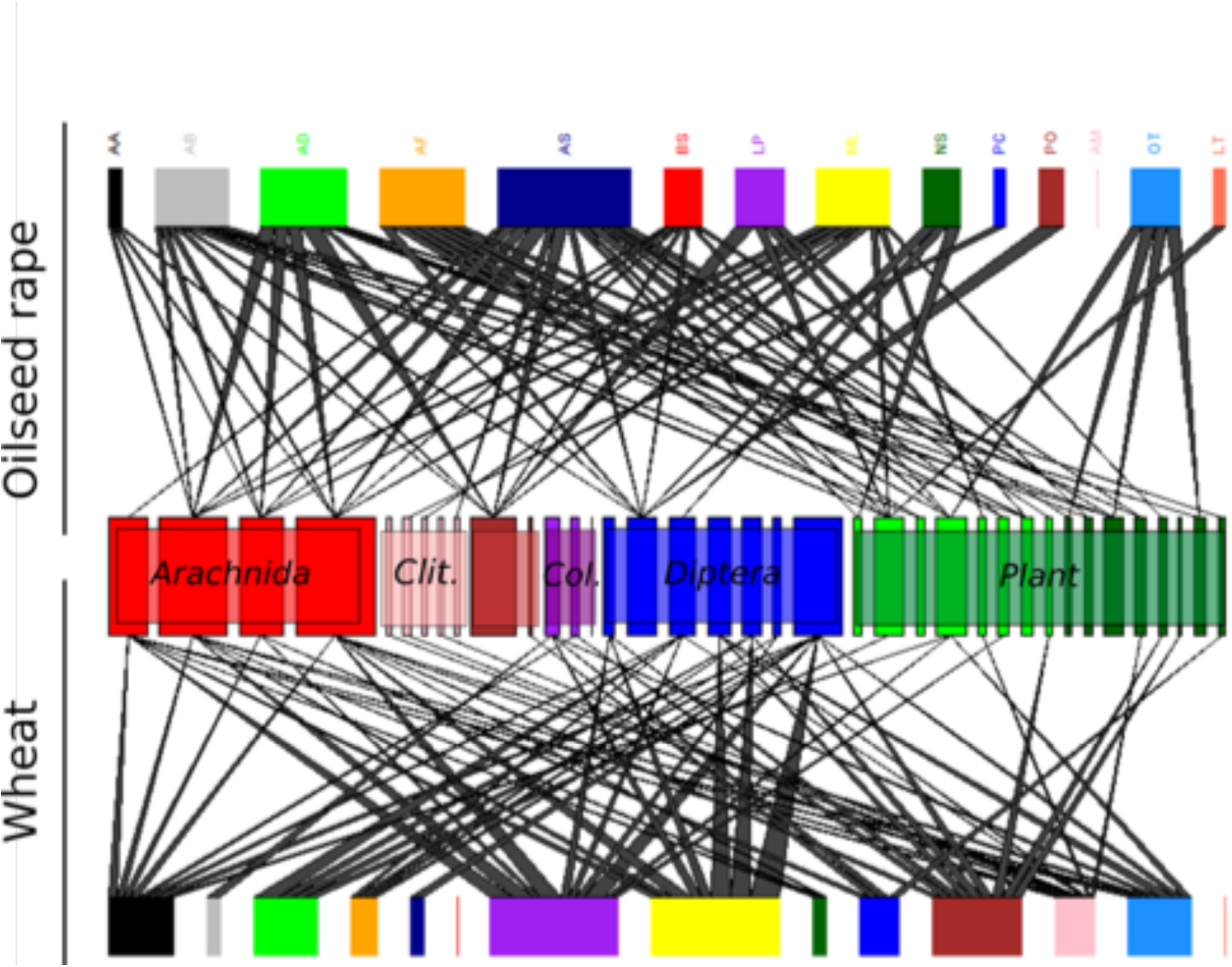
Bipartite food webs involving carabid beetles in their prey in oilseed rape and wheat fields. For a matter of clarity, trophic links between carabids and their resources are shown only for carabid individuals that were positive for at least on prey type. The width of segments represents the proportion of total links for the species. Abbreviations correspond to the fourteen carabid species that have showed positive for prey DNA (AA: *Amara aenea*; AB: *Anisodactylus binotatus*; AD: *Anchomenus dorsalis*; AF: *Asaphidion flavipes*; AM: *Agonum muëlleri*; AS: *Amara similata*; BS: *Brachinus sclopeta*; LP: *Loricera pilicornis*; ML: *Metallina lampros*; NS: *Nebria salina*; PC: *Poecilus cupreus*; PO: *Phyla obtusa*; OT: *Ocydromus tetracolus*; LT: *Laemostenus terricola*). The bars in the middle correspond to prey Operational Taxonomic Units (OTUs) retrieved from carabid gut contents. We have grouped the different OTUs into the following resource categiories: Arachnida (in red, 4 OTUs), Clitellata (in pink, 5 OTUs), Collembola (in brown, 2 OTUs), Coleoptera (in purple, 2 OTUs), Diptera (in blue, 7 OTUs), and Viridiplantae (in green, 16 OTUs).

**Figure 3.**
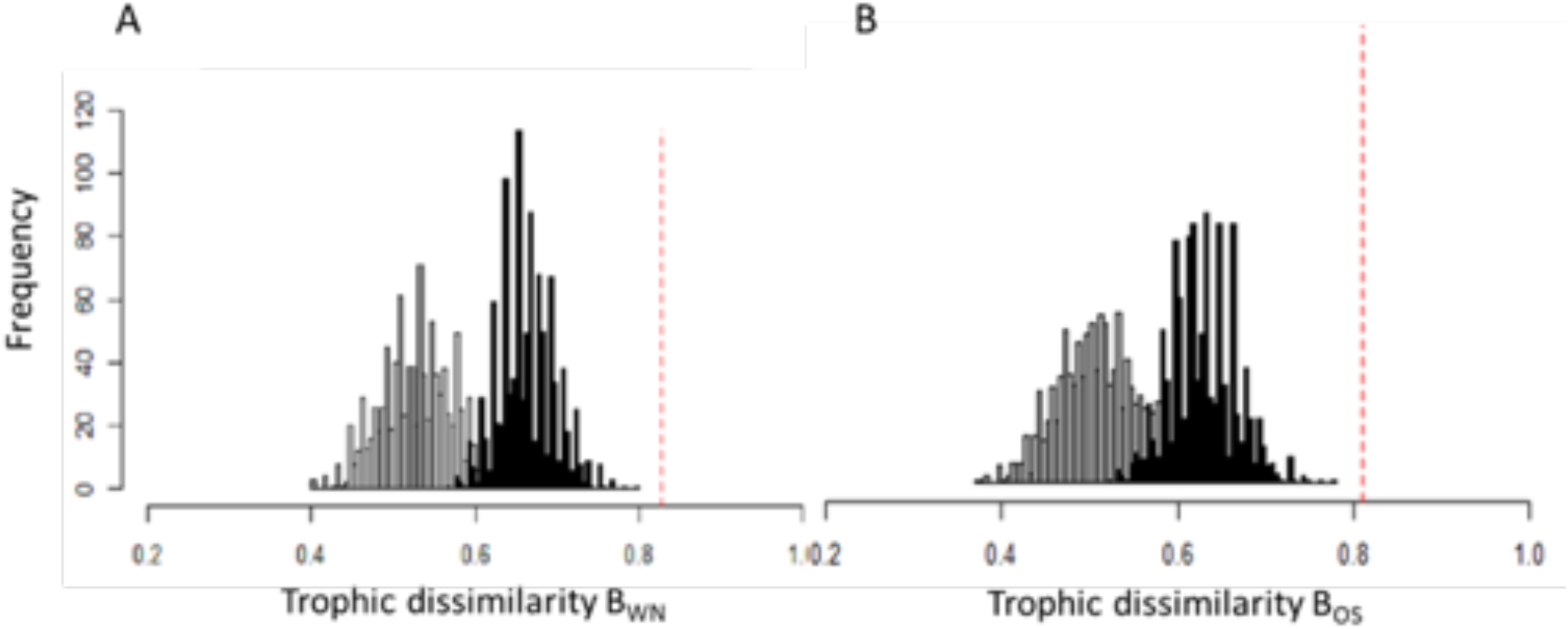
Distribution of trophic dissimilarity values between oilseed rape and wheat crops for (A) the whole networks (β_WN_) and (B) for the networks restrained to strong trophic links (β_OS_). Values correspond either to observed carabid food webs (red line) either to type 1 permutated carabid food webs (grey, individuals permutated between oilseed rape and wheat crops) or type 2 permutated carabid food webs (black, individuals of the same species permutated between oilseed rape and wheat crops). Results show that we have strong differences between oilseed rape and wheat food webs both at the scale of the whole network (β_WN_ > 0.8) and at the scale of species (β_OS_ > 0.8). Permutation simulations show that part of those differences resulted from the unbalanced distribution of positive carabids between the two crop types (grey for simulated data with individual permutation irrespective to their species – type 1) but also from differences in the identities of species positive for at least one prey in wheat and oilseed rape (i.e. effect of the community composition of the carabid testing positive for at least one prey between the two crop types, black-type 2 permutation). Note that network dissimilarity between crop types remained significantly higher than expected even when differences in community composition was taken into account.

**Fig. 4.**
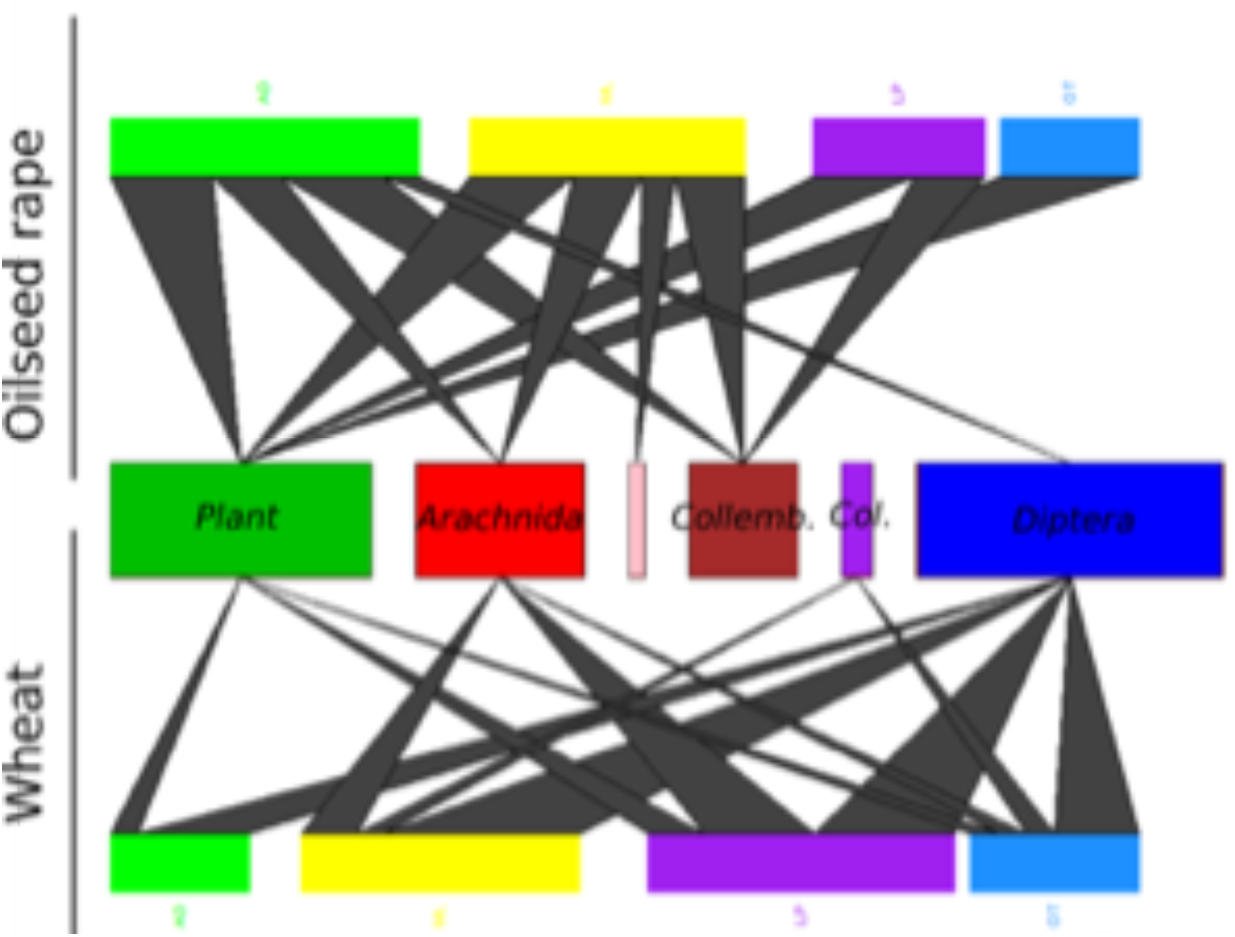
Bipartite carabid food webs representing only the strong trophic links in oilseed rape and wheat crops. Trophic networks in oilseed rape (up) and wheat (down) include only the carabid species for which at least 3 individuals where positive to at least one prey category within the two sampled crop types. From left to right prey categories correspond to Viridiplantae, Arachnida, Clitellata, Collembola, Coleoptera and Diptera. Carabid species abbreviations correspond to *Anchomenus dorsalis* (AD), *Metallina lampros* (ML), *Loricera pilicornis* (LP), *Ocydromus tetracolus* (OT).

## Discussion

In this study, we address the longstanding question of carabids’ contribution to biological control. Using a DNA metabarcoding approach, for the first time we explored the full diet spectrum and the biological control potential of a community of fourteen carabid species sampled within an intensively managed European agroecosystem. Our results confirm our main expectation that different carabid species exhibit different diets with various degrees of animal and plant pest consumption. We found within carabid gut contents a variety of animal and plant taxa including collembolans, insects, arachnids, earthworms and several plant families. All animal taxa are characteristic of the soil macro-fauna thus matching with the ground-dwelling foraging behaviour of carabid beetles. Among the animal and plant taxa detected within carabid gut contents, several included also potential pest species (dipterans, weed species). We also observed that carabids could prey upon beneficial organisms such as earthworms and spiders.

However, our results also suggest that there is no significant association between a carabid species and the consumption of a particular resource type. This finding confirms the generalist feeding behaviour of many of these predators and implies that the trophic choice in many common carabid species does not appear to be strongly constrained by their taxonomic identity. On the contrary, our results point towards the importance of environmental factors such as the field location and the crop type, which seem to drive carabid dietary choice in this case. We observed that at the same date, the same species could exhibit different diets according to the field location or the crop type where they have been collected. This is a novel finding because even if such dynamical changes in carabid feeding behaviour are known from previous studies, none of them have tempted to disentangle their effect from the effect of species identity while quantifying interactions at the community level. Previous studies have rather focused on single or a few predator species and / or have only quantified specific trophic linkages without considering the whole diet spectrum (e.g. King et al. 2010; Davey et al.2013). This is important, as one of the major challenges for agroecology today is the successful management of biodiversity that maximizes the delivery of ecosystem services such as biological control. Yet, managing biodiversity is challenging because of the difficulty of disentangling the effect of specific taxa from effects at the community level. Our approach suggests that carabid community composition and species identity do not seem to be good proxies for predicting biological control potential. Therefore, the presence of specific carabid taxa could not be used to infer levels of ecosystem services (biological control) or disservices (e.g. intraguild predation) as traditionally advocated. Rather, carabid beetles seem to adapt their feeding behaviour to their local environment. We observed higher consumption of dipteran prey in wheat crop, while more collembolans and plant material were consumed in oilseed rape fields. We speculate that the major determinant of this diet switch is the differences in resource availability between the two crop types (Smith et al. 2008). We did not directly quantify resources available for carabids but it is known, for instance, that oilseed rape fields are usually characterized by thick layers of litter and higher levels of moisture-conditions favouring soil macro-fauna such as collembolans and earthworms (cf Bohan et al. 2005; Smith et al. 2008). Previous studies have also shown that Collembola could be an important resource for several carabid species such as *Loricera pilicornis* (e.g. Hintzpeter & Bauer 1986) or *Anchomenus dorsalis* (Basedow 1994). The higher rates of dipteran DNA detection from carabids collected in wheat fields is harder to explain. With our DNA metabarcoding approach we were able to detect dipteran taxa such as Anthomyiidae and Cecidomyiidae, which could correspond to some abundant pest species attacking Brassicaceae crops in our study area, and whose overwintering pupal stages could still be present within wheat fields due to crop rotation. However, other dipteran taxa were also recovered and we currently lack knowledge about the spatial distribution and availability of this taxonomic group within the LTER “Armorique”. Furthermore, oilseed rape fields in our study area harboured more abundant and diversified communities of weed species, which could also explain the higher rates of plant DNA detection in this crop type. Generally speaking, such differences in diet preferences could be explained by significant spatio-temporal instability of agricultural landscapes – i.e. frequent changes in resource quality and distribution due to changes in the frequency and intensity of trea™ents would lead carabids to shift their diet. Such instability could be prominent even at finer scales with differences in field and crop characteristics driving differences in resource availability even between adjacent fields from the same crop type (cf Puech et al. 2014). This may explain why in some cases the location of the field from which carabid species were collected also explained a significant part of the variation in their observed diet.

It is important stress out here that carabid trophic choice could be also driven by other factors such as the number and the abundance of the other carabid species within the community, which we did not really took into account here. Whereas we optimized the sampling of the exact same species, we observed important variations in their relative abundances between the six different fields and the two crop types we sampled. Additionally, important differences in the overall community composition were observed with oilseed rape field harbouring in average higher abundances and higher number of carabid species, especially granivorous species such as *Amara* spp. or *Harpalus* spp. (unpublished data). It will be interesting in exploring these aspects by comparing for instance the changes in carabid diet composition and breadth across communities varying in their species richness.

It must be underlined however that all these observations are based on a relatively limited sample size resulting mainly from the overall low prey DNA detection rates, thus impeding the possibility of covering adequately the diet spectrum for each carabid species between fields and crop types. The low detection rates could be the consequence of both biological and methodological shortcomings due to our study model. Carabid beetles exhibit both high starvation levels at the population scale (Bilde & Toft 1998) and rapid digestion rates that could potentially result in high proportion of empty gut contents in field-collected individuals. Despite the short duration of trapping, digestion could have occurred during this period of starvation. On the other hand, methodological constraints such as the overwhelming amplification of carabid DNA could also lead to lower detection rates of prey DNA. Carabid DNA is likely preferentially amplified compared to the low-concentration, degraded DNA of the preys they have consumed. The competition during PCR may thus result in the absence of amplification of prey DNA. The much higher detection rates with the plant primers (where no competition between carabid and plant DNA is expected) seem to confirm this hypothesis. Interestingly, detection rates with the earthworm primers were low and we still observed some unexpected amplifications of carabid DNA. In this case, low detection rates most likely match consumption rates (i.e. carabids consumed little earthworm prey). The amplification of carabid DNA may in this case result from the lack of amplifiable target DNA within carabid guts. Such issues could be resolved by both increasing the number of PCR replicates as well as the sequencing depth in order to increase the probability of picking up the less abundant DNA molecules. Additionally, recently developed enrichment protocols aiming at limiting the collateral extraction and amplification of predator DNA could be used alongside (e.g. size selection of the target DNA, Krehenwinkel et al. 2016; Eitzinger et al. this issue). All these measures taken together should allow to increase the number of positively testing individuals in order to adequately estimate interaction frequencies at the species level.

It is also worth pointing here that we did not amplify any molluscan DNA with the MAV primers, while carabids within agroecosystems are reputed to frequently prey on slugs, which were also very abundant in our study area (especially in oilseed rape fields). This may be explained by the lower efficiency of these primers owing to the blocking primer we used alongside in order to prevent the amplification of human DNA (cf de Barba et al. 2014). As this blocker is not entirely specific and was also used in concentrations ten times higher compared to the MAV primers, it may have affected the amplification process. This would also explain the generally limited amplifications for arthropod DNA with this primer set as well.

Overall, although results presented here should be interpreted with caution, we claim that our study still brings insights matching findings from several previous studies that have used PCR-based methods for diet analysis, and which show that carabids exhibit very dynamical feeding behaviour (Bell et al. 2010; King et al. 2010; Staudacher et al. 2018). Our study seems to confirm for the first time this at much finer temporal (the date) and spatial scale (the field) and points out the importance of the crop type in determining feeding behaviour. The main advantage with our approach is that it allows to estimate carabid diet spectrum without any *a priori* and use this information to simultaneously quantify contributions to ecosystem services (biological control) and disservices (e.g. intraguild predation) at the community level. It adds to the increasing evidence that trophic choice of natural enemies within agroecosystems would be mainly driven by bottom-up processes related to agricultural practices and resource distribution / abundances (Lohaus et al. 2012; Tixier et al. 2013; Mollot et al. 2014; Poeydebat et al. 2017) and we encourage future studies using DNA metabarcoding diet analysis in order to further support these findings.

## Data accessibility

Carabid-prey data matrix used for statistical analyses as well as the list of specimens sequenced for the COI prey reference database are available in the Supplementary section. Reference sequences will be deposited on GenBank while DNA metabarcoding sequence data and bioinformatic pipeline details will be deposited on FigShare after the acceptance of the manuscript.

## Acknowledgements

We thank all the field assistants and students from the UMR IGEPP who occasionally gave a hand with the collection and the sorting of the carabid beetles: Frédéric Hamelin, Nolwenn Génuit, MaeCl Dugué, Kévin Tougeron, Théo Vantsteenkeste, Frédérique Mahéo, Lucie Mieuzet, Nathalie Leterme, Jean-FrancCois Le Gallic, Bernard Chaubet and Sarah Polin. We also thank the LTER area ‘Armorique’ for providing infrastructures as well as the network of farmers that kindly allowed us to sample in their fields. Finally, we warmly acknowledge all the help we received from colleagues during the constitution of the prey reference database: Jean-Yves Rasplus, Astrid Cruaud, Jean-Claude Streito, Isabelle Meusnier, Maxime Galan and Gwenaelle Genson (UMR CBGP, INRA Montpellier); Julien Pétillon and Armelle Ansart (UMR ECOBIO, Université de Rennes 1); Jean-Luc Roger (SAD Paysage, INRA, Rennes); Marylin Roncoroni (CEBC Chizé); Nadya I. Nikolova (Biodiversity Institute of Ontario, Canada).

The authors of this study received funding from the Agence Nationale de la Recherche (‘Landscaphid’ project, ANR-09-STRA-05) and INRA depar™ent “Santé des Plantes et Environnement”. Stefaniya Kamenova was partially funded by the Région Poitou-Charentes (France) during her PhD thesis.

## Author’s contributions

SK, VB and MP designed the study, SK and MP collected the samples, SK carried out the molecular and bioinformatic analyses, EC carried out all statistical analyses with input from MP. SK wrote the paper with input from EC. MP and VB commented the final version of the manuscript.

## Supplementary material

### Appendix S1. Prey sequence reference database for the COI gene

**Table S1.**
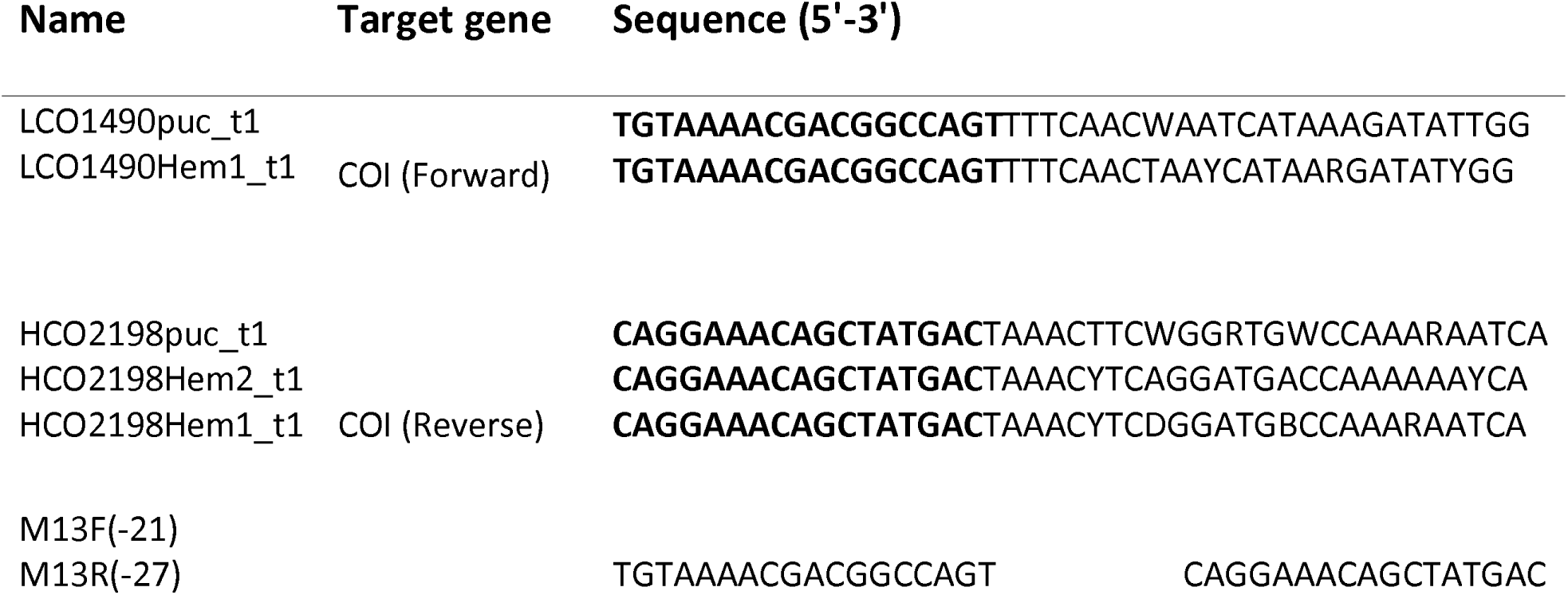
COI primer sequences used for the molecular referencing of the major prey groups encountered in our study area and that could be consumed by carabid beetles. The M13-tailed COI primer cocktail is prepared by pooling an equal volume of 10 uM of the five primers forward and reverse primers listed here. Characters in bold indicate the universal M13 tails. These tails play no role in amplification of the target but are used for generating cycle sequence products.

### Appendix S2. Sensibility analysis for GLM models of resources consumption by carabids individuals

We implemented a GLM models selection by selecting different carabid individuals. For this, we first included only species in common between the two crop types (therefore excluding *A. muelleri, B. bullatus, B. sclopeta*, and *L. terricola*), see table S2. Second, we removed one species at a time (Fig. S1). Results were particularly consistent for the Coleoptera, Arachnida and Plant prey categories, with the same significance levels and very similar coefficients between the two GLMs (Table S2, Fig. S1). For Diptera, only the effect of the field location was marginally significant and became non-significant when removing *A. aenea, A. binotatus, A. flavipes* or *A. similata* carabid species (Table S2). Earthworms and Collembola prey categories were also sensible to individual species removing. Results for earthworms were always non-significant when removing any of the carabid species, while Collembola was only sensitive to the removal of the *P. cupreus* species.

**Table S2. Best GLMs of resources consumption.** Generalized Linear models of OTUs presence / absence in carabids gut contents as a function of the carabid species, the crop type and the field location. We ran the GLMs by only retaining the species that were represented by at least 3 individuals testing positive for at least one prey category in the two crop types.

**Figure S1.** Coefficient of the best Generalized Linear models of OTUs detection in carabids gut contents as a function of the carabid species, the crop type and the field location. “All_sp” model is the same as presented in the manuscript, including all the fourteen carabid species. Thirteen model selections have been carried out in the same way, except that one species at a time was removed (for instance in the “no_AA” model, the best model was obtained when individual from the *Amara aenea* species has been removed). (AA: *Amara aenea*; AB: *Anisodactylus binotatus*; AD: *Anchomenus dorsalis*; AF: *Asaphidion flavipes*; AM: *Agonum muëlleri*; AS: *Amara similata*; BS: *Brachinus sclopeta*; LP: *Loricera pilicornis*; ML: *Metallina lampros*; NS: *Nebria salina*; PC: *Poecilus cupreus*; PO: *Phyla obtusa*; OT: *Ocydromus tetracolus*; LT: *Laemostenus terricola*).

